# Potential role of extracellular ATP released by bacteria in bladder infection and contractility

**DOI:** 10.1101/538868

**Authors:** Behnam Abbasian, Aidan Shair, David B. O’Gorman, Ana M. Pena-Diaz, Kathleen Engelbrecht, David W. Koenig, Gregor Reid, Jeremy P. Burton

## Abstract

Urgency urinary incontinence (UUI), the result of conditions such as overactive bladder (OAB), could potentially be influenced by both commensal and urinary tract infection-associated bacteria. The sensing of bladder filling involves interplay between various parts of the nervous system eventually resulting in contraction of the detrusor muscle during micturition. Here we model host responses to various urogenital bacteria, firstly by using urothelial bladder cell lines and then with myofibroblast contraction assays. To measure responses, we examined calcium influx, gene expression and alpha smooth muscle actin deposition assays. We found that organisms such as *Escherichia coli* and *Gardnerella vaginalis* strongly induced calcium influx and contraction, whereas, *Lactobacillus crispatus* and *L. gasseri* did not induce this response. Additionally, supernatants from lactobacilli impeded influx-and contraction induced by the uropathogens. Upon further investigation of factors associated with the purinergic signaling pathways, we found that influx and contraction of cells correlated to the amount of extracellular ATP produced by *E. coli*. Certain lactobacilli appear to mitigate this response by utilizing extracellular ATP or producing inhibitory compounds which can act as a receptor agonist or calcium channel blocker. These findings suggest that members of the urinary microbiota may be influencing UUI.

## IMPORTANCE

The ability of the uropathogenic bacteria to release significant amounts of ATP as an excitatory compound and possible virulence factor to stimulate various signaling pathways can have profound effects on the urothelium, perhaps extending to the vagina. This may be countered by the ability of certain commensal urinary microbiota constituents, such as lactobacilli. The clinical implications are to better understand the impact of antimicrobial therapy on the urinary microbiota and to develop a more targeted approach to enhance the commensal bacteria and reduce ATP release by pathogens.

## INTRODUCTION

Patients who suffer from overactive bladder syndrome (OAB) or urgency urinary incontinence (UUI) usually experience the sensation to urinate whether the bladder is full or not. While there are many factors involved, ultimately it is the contraction of bladder smooth muscle cells which invokes urination [1, 2]. Storage and voiding of urine are controlled by sympathetic and parasympathetic pathways [1,3], via the adrenergic and cholinergic systems. It has been speculated that neurotransmitters with different effects and potentially originating from bacteria, may play major roles in bladder function [4, 5].

The discovery of a urinary microbiota has shown that diversity differs between healthy people and patients with neurogenic bladder dysfunction, interstitial cystitis, UUI and sexually transmitted infections [6, 7, 8]. The microbial diversity in women with UUI may be associated with severity of the condition [9]. The genus *Lactobacillus* has been found more frequently in healthy subjects compared to patients with UUI (60% versus 43%), while *Gardnerella* was more abundant in patients (26% versus 12% in controls) [9]. Interestingly, in some studies, *L. gasseri* is considerably more prevalent in UUI patients than *L. crispatus* raising questions about how different species adapt to the bladder [9].

It may seem difficult to envisage how the detrusor muscle which controls micturition could be affected by bacteria present at the urothelial layer. Yet, the urothelium is only 3-5mm thick, and uropathogens have been shown to damage and invade this layer [10]. Urothelial cells communicate with the sub-urethral tissue in the lamina propria, which contains nerve fibers and smooth muscle cells, by releasing excitatory compounds such as ATP [10,12]. Bacterial compounds could induce urothelial cells to release excitatory compounds into the sub-urethral space, thereby inducing smooth muscle contraction and voiding. We hypothesize that bacteria produce, release and potentially sequester compounds, such as ATP that play a role in UUI pathogenesis and that commensal bacteria may be beneficial to prevent detrusor muscle contractions.

Here we explore interactions of uropathogenic bacteria and commensal lactobacilli to affect the physiology of bladder cells in culture and to release ATP to stimulate calcium influx and contraction of myofibroblasts.

## RESULTS

### Ca^2+^ influx of uroepithelial cell by bacterial supernatants

The supernatant of *E. coli 1A2* was able to induce the influx of Ca^2^+ [Fig 1A]. Unlike previous reports [13], LPS did not rapidly stimulate the influx of calcium in our model [Fig 1A]. Supernatants from *E. coli* increased the levels of Ca^2+^ influx before plateauing at the 4-hour time point. In contrast, *E. faecalis* 33186 supernatant did not significantly increase the levels of Ca^2+^ influx until the 5-hour time point [Fig 1B and C].

**Fig 1:**
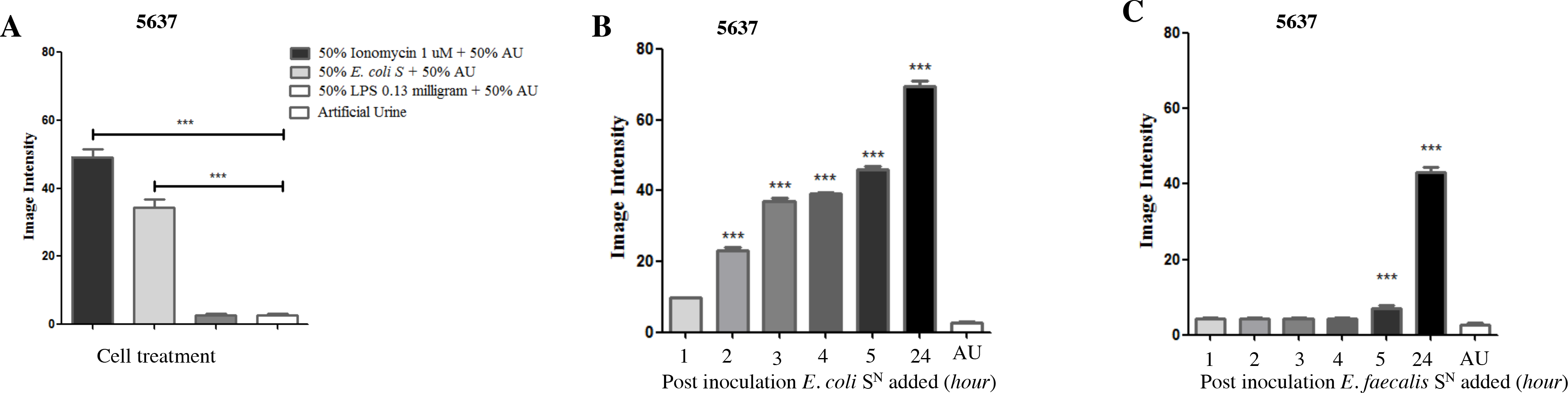
Effect of *E. coli* and *E. faecalis* on the stimulation of Ca2+ influx in 5637 uroepithelial cells. Bacterial supernatant (S^N^) was added to the 5637 cells (**A**), S^N^ from either *E. coli* (**B**) or *E. faecalis* (**C**) were taken from hourly growth at 1, 2, 3, 4, 5, 24 hours and artificial urine (AU) and tested for their ability to induce Ca^2+^ influx in the 5637 cells. Each bar represents the total average image intensity 60 seconds after treatment of a duplicate sample. Statistical significance was determined using Tukey’s test, p≤0.05.

### *Lactobacillus crispatus* ATCC 33820 and *Lactobacillus gasseri* KE-1 supernatants reduce Ca^2+^ influx caused by *Escherichia coli*

The addition of *L. crispatus* and *L. gasseri* supernatants mitigated the effects on calcium influx caused by *E. coli* supernatant up to 50% [Fig 2A-D].

**Fig 2:**
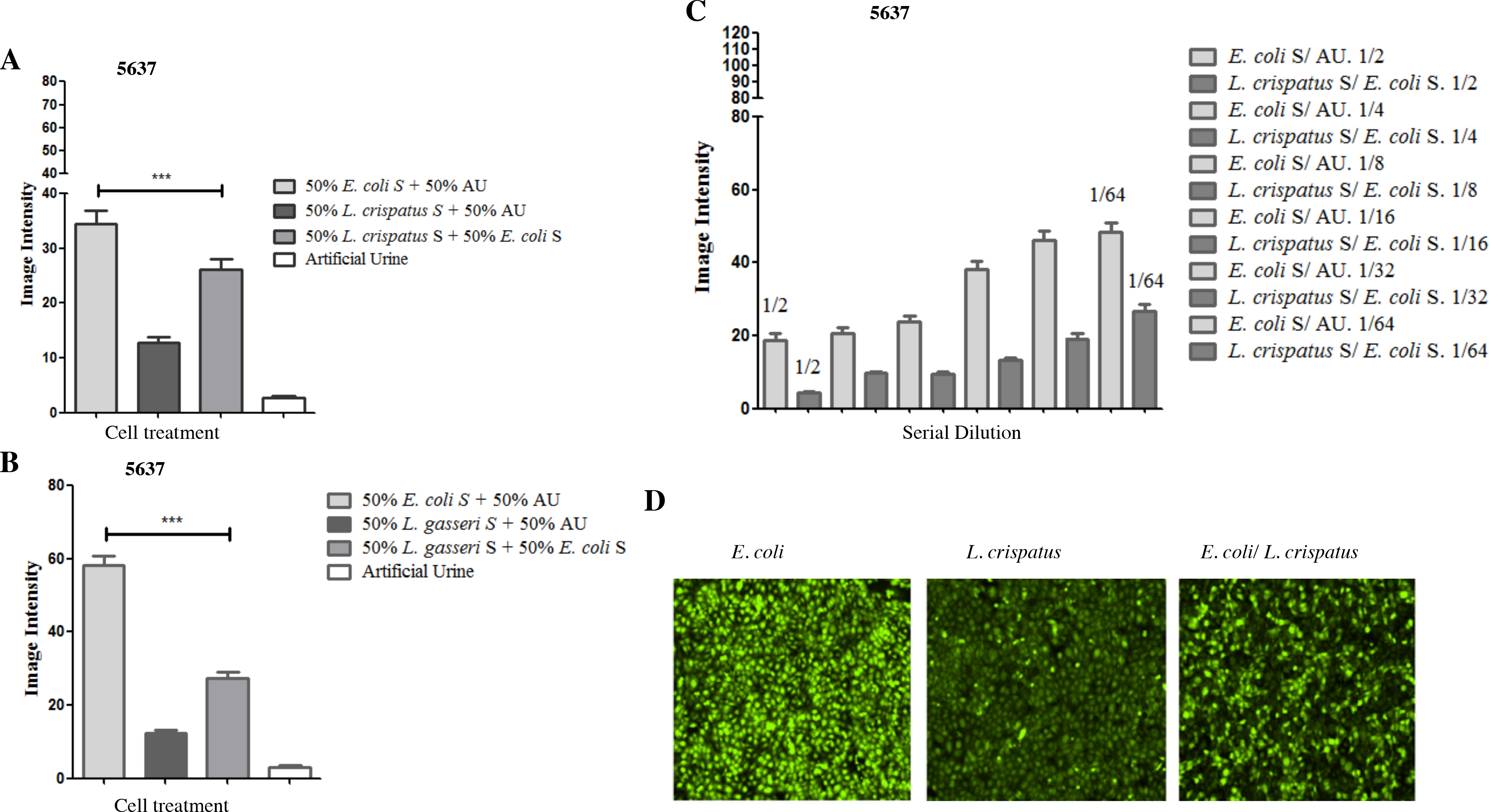
Effect of *L. crispatus* and *L. gasseri* on Ca^2+^ influx caused by UPEC supernatant. Treatments were added as either *E. coli S*^*N*^ and AU or *L. crispatus S*^*N*^ and AU, or *E. coli S*^*N*^ + *L. crispatus* S^N^ (**A**), or *E. coli S*^*N*^ and AU or *L. gasseri* S^N^ and AU, or *E. coli S*^*N*^ + *L. gasseri* S^N^ (**B**), or a six-fold serial dilutions of the *L. crispatus* S^N^ in the *E. coli* S^N^ (**C**), image examples of calcium influx caused by *E. coli, L. crispatus* and a mixture of S^N^ from the two bacteria (**D**). Each point represents the average image intensity of a timepoint. Statistical significance was determined using Tukey’s test, p≤0.05.

### Quantification of bacterial extracellular ATP

A luminescent assay was used to quantify the amount of extracellular ATP released by bacterial supernatants. The *E. coli*, *L. crispatus* and *L. gasseri* supernatants from the overnight culture contained 0.098 ± 0.008 uM, 0.024 ± 0.003 uM and 0.024 ± 0.001 uM ATP, respectively, which was significantly higher than AU 0.0067 ± 0.0011 uM [Fig 3A]. In addition, supernatants of urinary microbiota constituents, *G. vaginalis* ATCC 14018 and *L. vaginalis* NCFB 2810 contained 1.30 ± 0.14 uM and 0.314 ± 0.023 uM ATP, respectively, which was significantly greater than the AU control of 0.0067 ± 0.0011 uM [Fig 3A and 3B].

**Fig 3:**
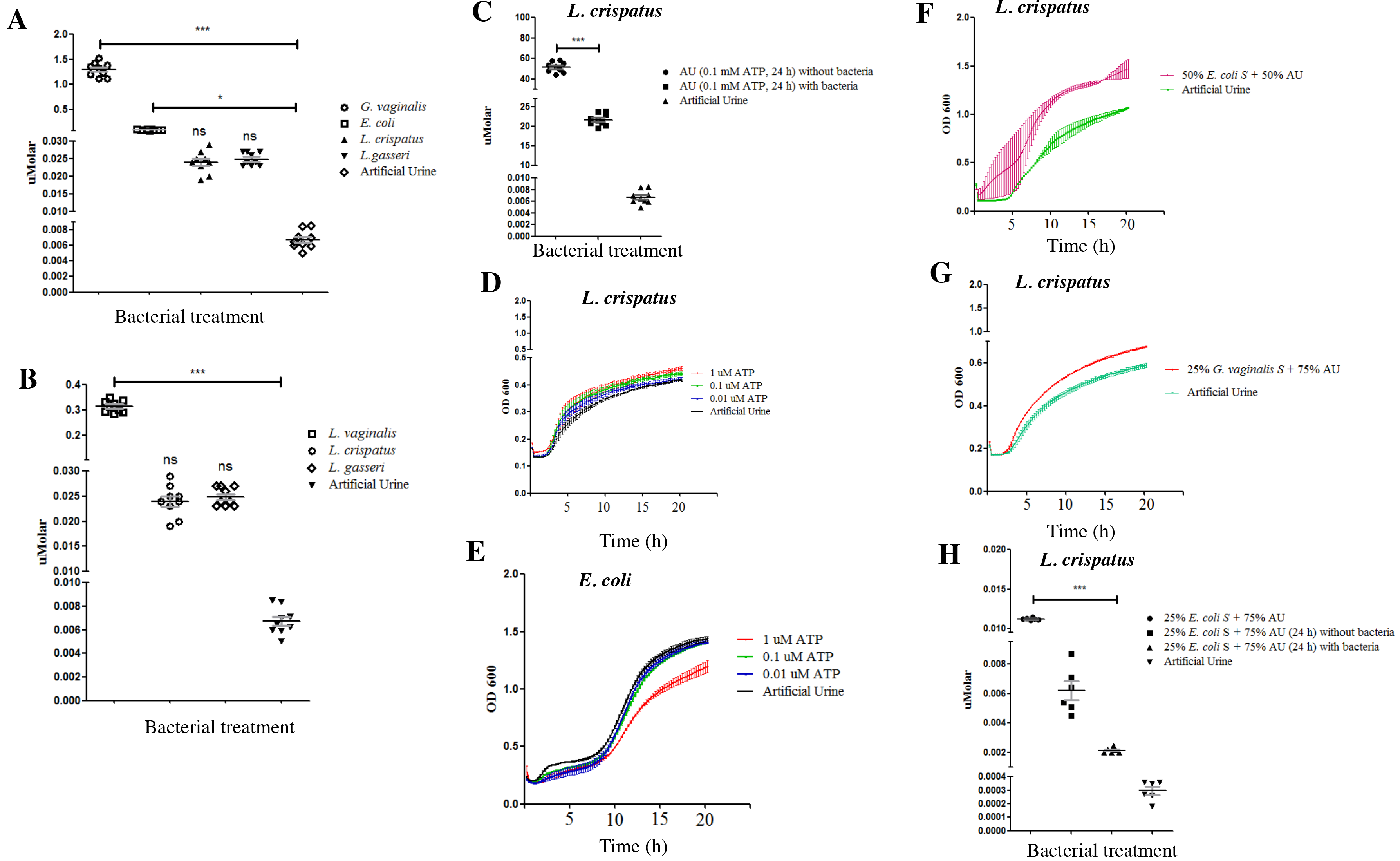
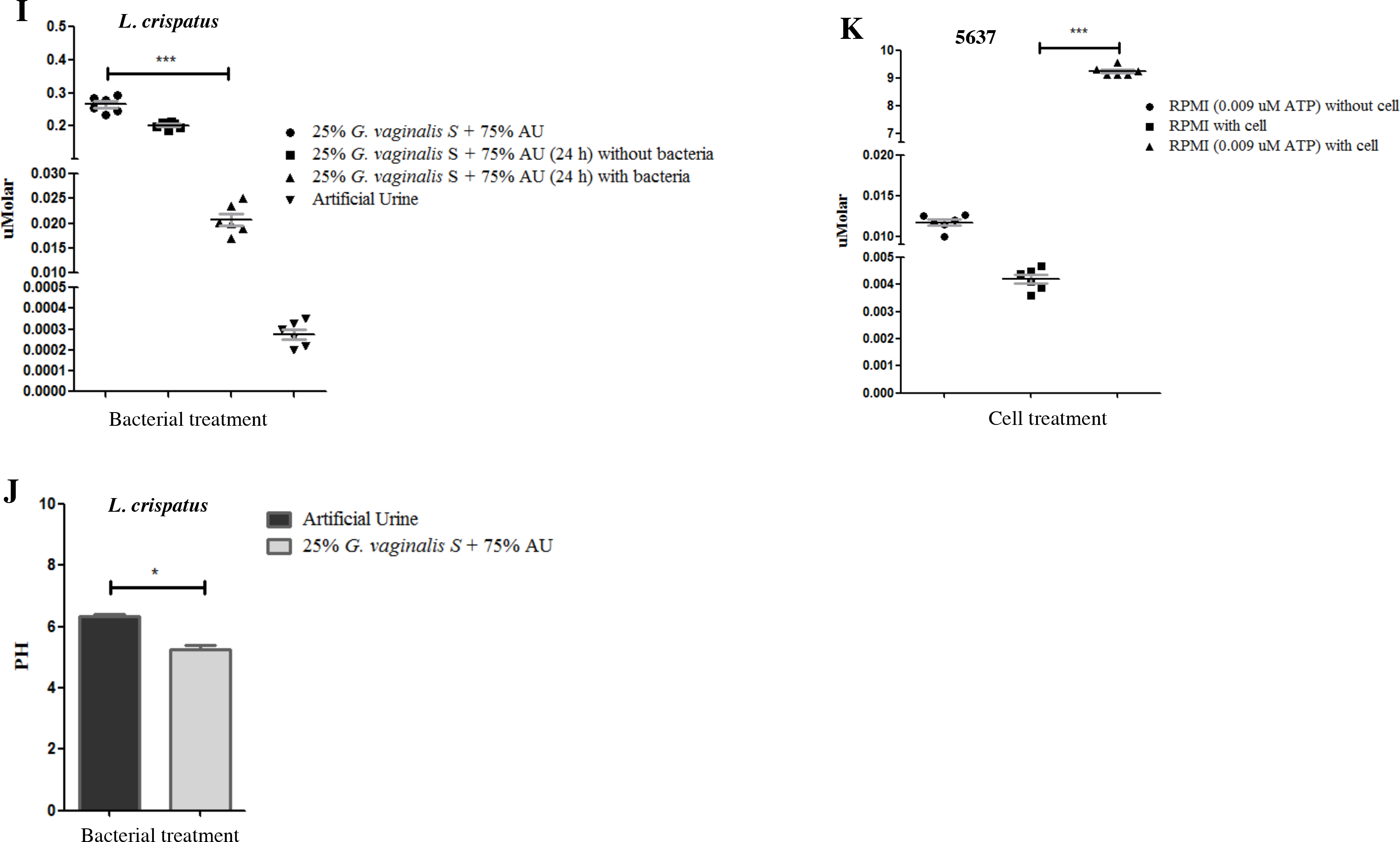
Release of extracellular ATP by bacteria. *E. coli*, *L. crispatus*, *L. gasseri*, *G. vaginalis and L. vaginalis* S^N^ were collected from overnight cultures grown in AU and measured for ATP (**A and B**). Statistical significance was determined using Tukey’s test, p≤0.05. *L. crispatus* was grown in AU supplemented with 0.1 millimolar of ATP overnight, the amount of ATP was evaluated by luminometer (**C**). Growth of *L. crispatus* and *E. coli* was measured in the presence of different concentrations of ATP in AU (**D and E**), additionally for *L. crispatus* supplemented with *E. coli* or *G. vaginalis* supernatants (**F and G**). The ability of *L. crispatus* to reduce the amount of ATP in AU supplemented with 25% of *E. coli* S^N^ (**H**), and 25% *G. vaginalis* S^N^ individually, was also examined (**I**). The ability of 25% of *G. vaginalis* to induce *L. crispatus* to reduce pH was also examined (**J**). RPMI was supplemented with small quantities of ATP and incubated for two minutes. ATP was measured before and after the addition of the ATP (**K**).

The amount of ATP remaining when *L. crispatus* was grown in AU supplemented with 0.1 mM ATP for 24 hours was 21.67 ± 1.51 uM, less than half the control (51.56 ± 5.06 uM) (P≤0.0001) [Fig 3C]. To investigate ATP reduction, *L. crispatus* was cultured in AU supplemented with different concentrations of ATP, as well as in AU supplemented with 50% *E. coli* supernatant, and 25% *G. vaginalis* supernatant, as potential natural sources of ATP. The growth of *L. crispatus* was increased by increasing ATP concentration, including that emanating from the *E. coli* and *G. vaginalis* supernatants [Fig 3D, F-I]. *Lactobacillus crispatus* also reduced the amount of ATP after overnight culture in AU supplemented with 25% of *E. coli* supernatant, and 25% of *G. vaginalis* supernatant individually. Supplementing *E. coli* with ATP had a somewhat inhibitory effect on its growth [Fig 3E]. In the presence of ATP or supernatant from *G. vaginalis*, the pH of *L. crispatus*, became further reduced (Fig 3J).

### Urothelial cells were forced to release ATP

The urothelial cell media contained 0.0042 ± 0.00040 uM ATP, and after treatment with 0.009 uM ATP for 2 minutes, the ATP released by the urothelial cells increased to 9.237 ± 0.172 uM [Fig 3K].

### The effects of sub-therapeutic ciprofloxacin on *E. coli* to release ATP

After culturing bacteria with different concentrations of ciprofloxacin from 10 ug/mL to 0.031 ug/mL, the minimum inhibitory concentration (MIC) of the antibiotic against *E. coli* was 1 to 1.5 ug/mL. Using MIC concentrations of ciprofloxacin below the MIC of 0.25, 0.125, 0.0625 ug/mL induced *E. coli* to release more ATP up to 0.0247 ± 0.0015 uM [Fig 4].

**Fig 4:**
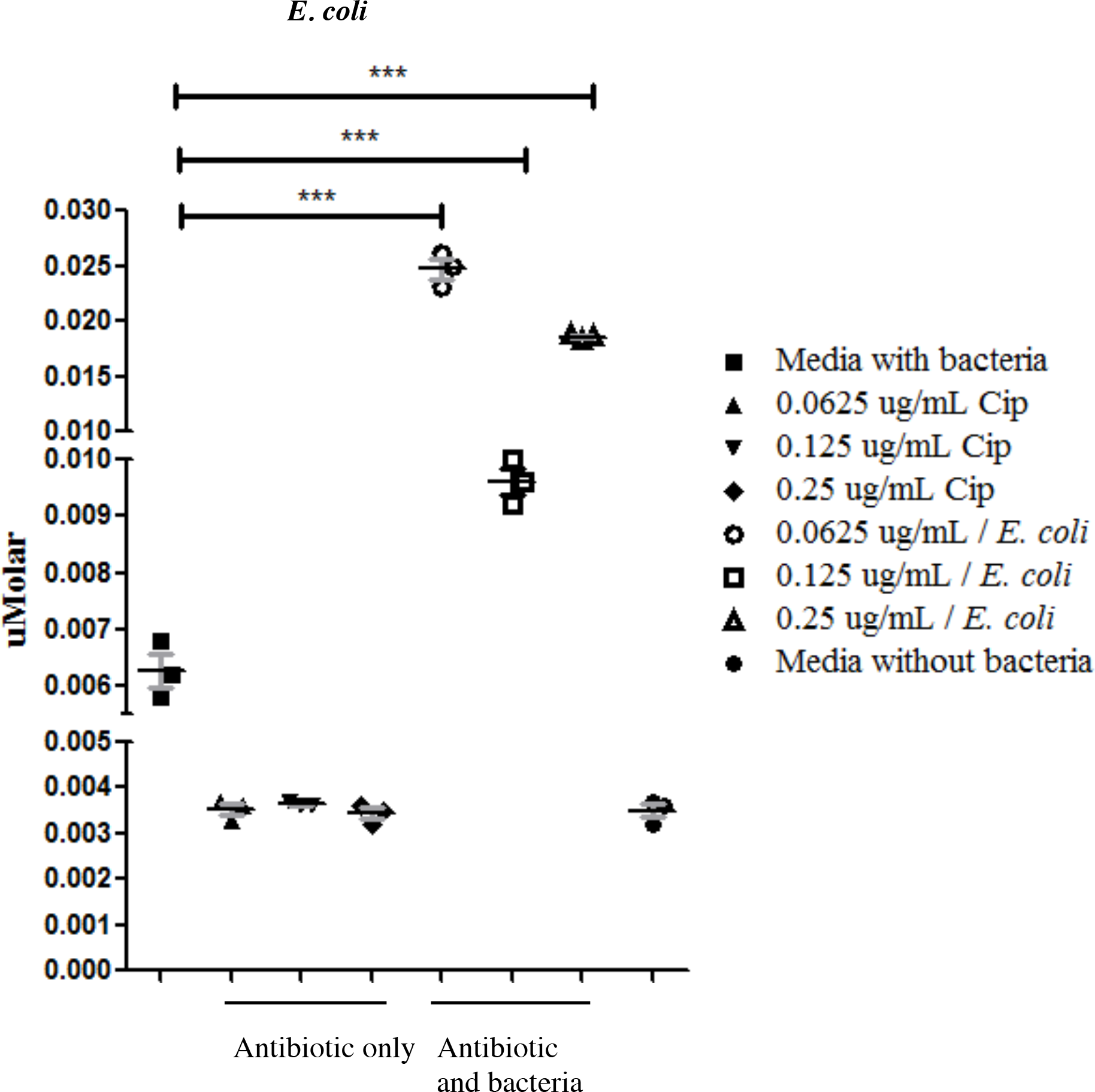
Release of ATP by *E. coli* in the presence of ciprofloxacin. This was to determine the minimum inhibitory and subtherapeutic concentrations of exposure to this antibiotic. *E. coli* were grown overnight culture in various sub-MIC concentrations of ciprofloxacin and releases significant quantities of ATP at different sub-MIC antibiotic concentrations.

### Expression of *MAOA* and *MAOB* in the 5637 cells exposed to bacterial supernatants

ATP by increasing the level of intracellular calcium can cause mitochondrial dysfunction. Gene expression for mitochondrial enzymes, monoamine oxidase A and B was measured because of their potential ability to degrade neurotransmitters such as serotonin. The *E. coli* supernatant (−1.27 ± 0.0041 fold change), as well as *L. crispatus* supernatant (1.218 ± 0.0020 fold change) had no effect on *MAOA* gene expression [Fig 5A]. The *E. coli* had no effect on *MAOB* gene expression, whereas, *L. crispatus* upregulated its expression by 44-fold [Fig 5B].

**Fig 5:**
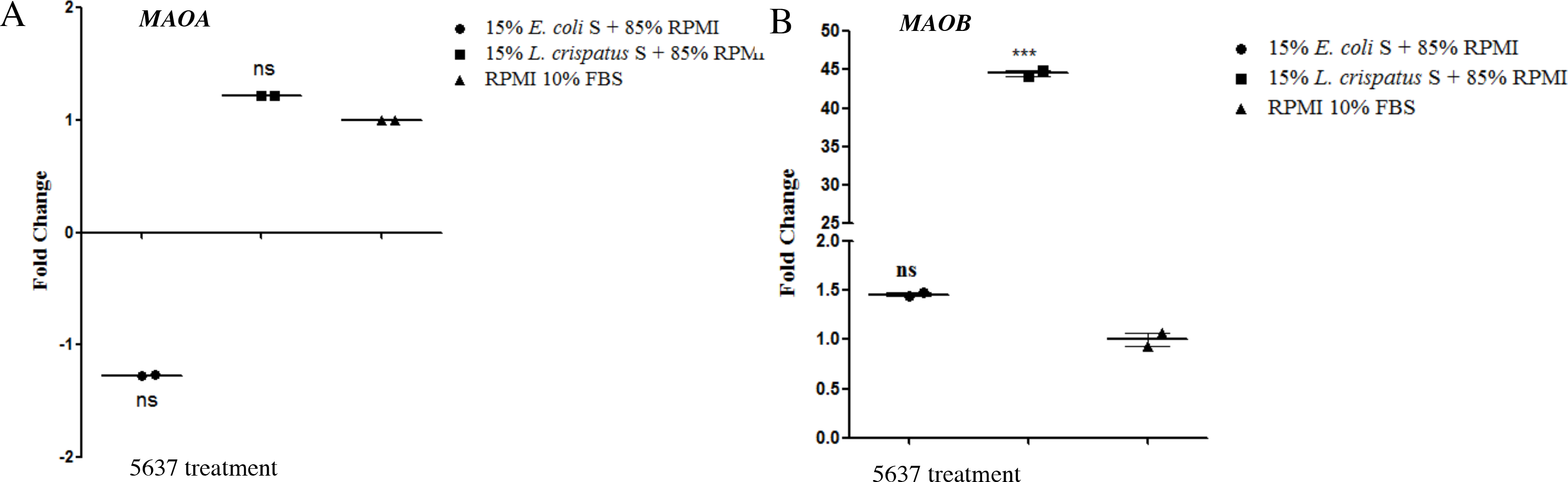
Fold change of transcript expression of monoamine oxidases. Supernatants from both pure cultures and mixtures of *E. coli* and *L. crispatus* S^N^ were added to 5637 cell cultures for 3 hours. Expression of genes encoding monamine oxidases (*MAOA/ MAOB*) were measured by quantitative PCR relative to GAPDH (**A and B**).

### Investigation the effect of GABA on Ca^2+^ influx caused by ATP and bacterial supernatant

The neurotransmitter γ-aminobutyric acid (GABA) was found to reduce the stimulation of calcium influx caused by ATP [Fig 6A] and inhibit the stimulation of calcium influx caused by *E. coli* supernatant [Fig 6B].

**Fig 6:**
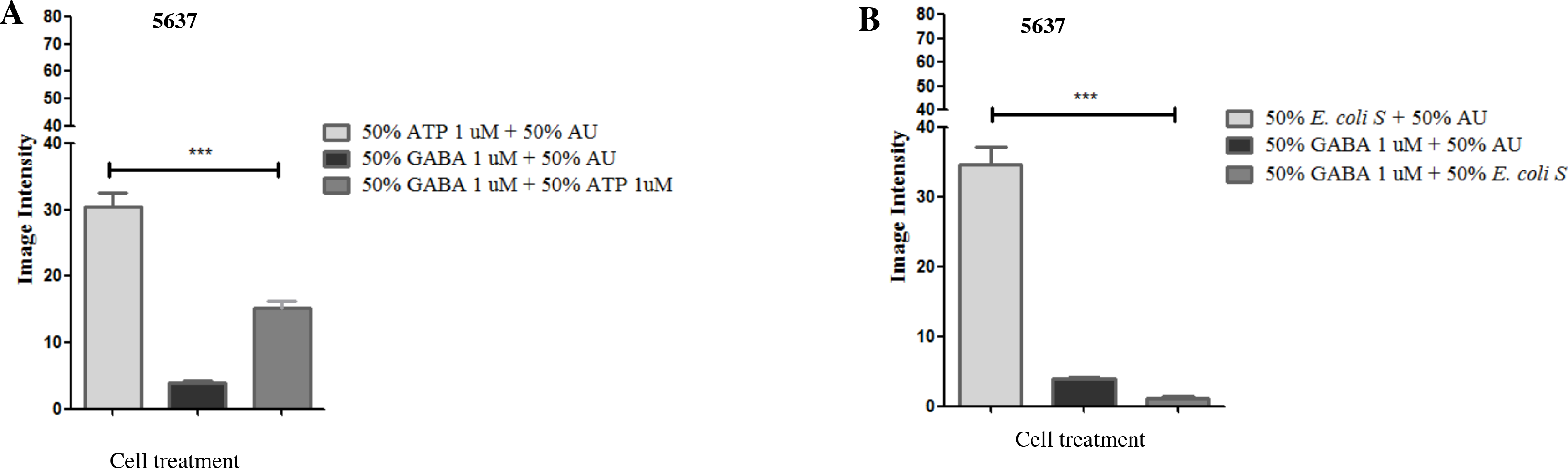
Effect of GABA and ATP on Ca^2+^ influx in 5637 cells. To evaluate the ability of GABA to inhibit the stimulation of calcium influx caused by ATP, AU containing 1 uM GABA was mixed with 1 uM ATP in AU (**A**) Similarly, to test the ability of GABA to reduce the stimulation of calcium influx caused by bacterial supernatant, GABA was mixed with *E. coli* supernatant (**B**). Statistical significance was determined using Tukey’s test, p≤0.05.

### Myofibroblast contraction assay

A collagen contraction assay using primary myofibroblast cells seeded inside a collagen matrix was tested against bacterial products as an *in vitro* model of smooth muscle contraction. Supernatant from cultures of *E. coli* were able to induce the greatest amount of contraction (72.67% ± 0.87) in the myofibroblast cell line after 24 hours and this reduced when *L. crispatus* or *L. gasseri* supernatants were added (48.56% ± 1.68, 29.82% ± 0.023, respectively) [Fig 7A, B and 7C]. Pure ATP caused contraction of myofibroblasts in the first hour (30.30% ± 3.25) and continued for 24 hours (60.73 % ± 1.49) [Fig 7D]. While, GABA did not cause contraction in the myofibroblast assay, it inhibited contraction caused by *E. coli* [Fig 7E]. Previous reports [13] suggest that the contraction maybe caused by *E. coli* was due to LPS. However, after five hours of exposure of LPS to the myofibroblasts, contraction was approximately half that induced by ATP (27.26% ± 1.05 versus 46.19 ± 1.78%). [Fig 7F].

**Fig 7:**
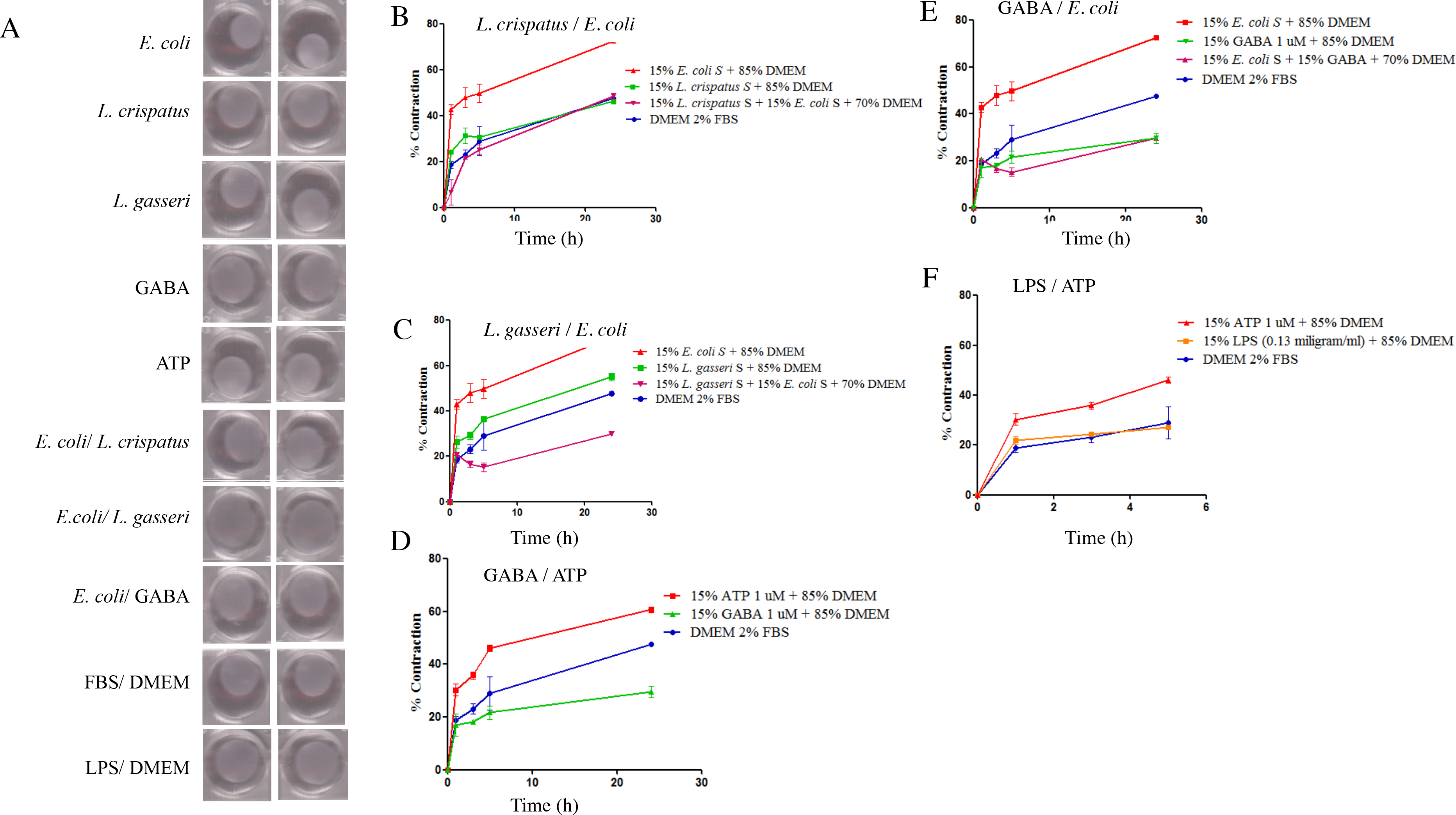
Contraction of a myofibroblast populated collagen matrix by bacterial supernatants. Bacterial supernatants from *E. coli, L. crispatus* and *L. gasseri* were added to a myofibroblast populated collagen matrix, both from pure culture and mixtures with DMEM with 2% FBS. In addition, GABA, ATP and LPS were included as controls. Contraction over time for these treatments is also shown (**B-F)**.

### Immunocytochemistry for intracellular alpha smooth muscle actin (α-SMA) and induction of TNF by bacteria

To further confirm myofibroblast contractive abilities in the presence of bacterial compounds, the effect on alpha smooth muscle actin was assessed. The *E. coli* supernatant increased the intracellular image intensity (56.43 ± 2.86) which is related to the alpha smooth muscle actin [Fig 8A and B] and this was reduced by *L. crispatus* (13.4 ± 1.45). The *E. coli* supernatant did not increase the expression of the *ACTA2* (−1.037 ± 0.023 fold change) [Fig 8C]. *Lactobacillus crispatus* also downregulated the level of *ACTA2* gene expression (−1.7 ± 0.029 fold change) [Fig 8C] Thus, the ability of *E. coli* to increase the intracellular image intensity could potentially be based on alpha smooth muscle cells contraction, and the ability of *L. crispatus* to reduce the intracellular image intensity could potentially be based on alpha smooth muscle cells relaxation.

**Fig 8:**
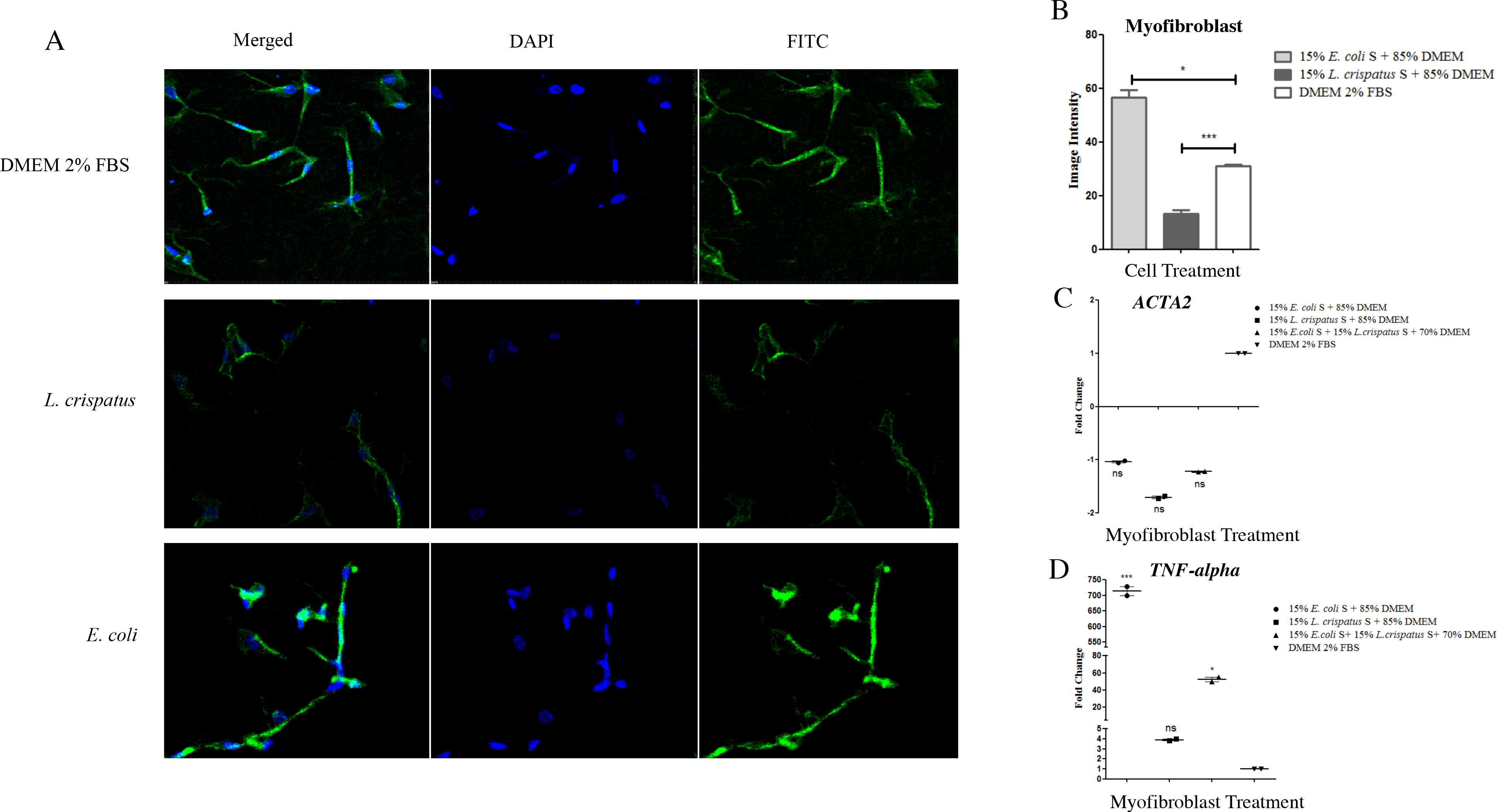
Confocal fluorescence microscopy analysis. Bacterial supernatants from *E. coli, L. crispatus* and in combination were co-cultured with myofibroblasts for 1 hour (**A and B**). The image intensity was measured by confocal microscopy with DAPI and FITC to show staining of α-smooth muscle actin. Bacteria supernatants identical to A, were added to the myofibroblasts that were grown in the collagen matrix and then incubated at 37°C with 5% CO_2_ for 3 hours and tested for gene expression of alpha smooth muscle actin (*ACTA2*) and TNF (**C and D**).

To determine if sustained activation of the calcium channel promoted apoptosis by bacterial components, *TNF* was measured as an indicator. the *E. coli* caused more than 700-fold upregulation (714 ± 19.91) of *TNF* [Fig 8D], whereas, exposure to *L. crispatus* only resulted in three-fold increase (3.9 ±0.115). When *E. coli* and *L. crispatus* supernatants were mixed and applied to the assay, the expression of *TNF* induced by the *E. coli* was strongly mitigated (52.38 ± 3.98) [Fig 8D].

## Discussion

Here we show that uropathogenic *E. coli* can release ATP into artificial urine and cause the influx of calcium [Fig 3A and 2A]. The ability to stimulate the uroepithelium could impact the sub-urethral space and smooth muscle cells and may directly affect the contractility of the bladder [14]. Studies using myofibroblasts showed that the *E. coli* supernatant induced high levels of collagen matrix contraction after 24 hours [Fig 7A/B].

Intracellular calcium has many roles inside the cell and regulates important mechanisms such as gene expression, metabolism, and proliferation [15]. This influx can be rapidly induced in the presence of ATP and has been previously shown to be produced extracellularly by *E. coli*, *Salmonella*, *Acinetobacter*, *Pseudomonas*, *Klebsiella* and *Staphylococcus in vitro* [12]. In patients with urinary infections, antibiotics are often administered. This reduces the number of bacteria in the lumen where they are exposed to therapeutic concentrations of the antibiotic. However, bacteria can also be embedded intracellularly in the urothelial cells, where only sub-therapeutic concentration of antibiotics may reach [16]. We have now shown that subtherapeutic exposure to ciprofloxacin induced *E. coli* to release more ATP [Fig 4], which could increase bladder contractions.

The role that the urinary microbiota of incontinence patients might have in uncontrolled voiding is supported by the finding of an abundant member of the microbiota, *G. vaginalis* releasing comparatively large amounts of ATP (1.30±0.14 uM) [Fig3A]. If these amounts were produced *in vivo* they would likely cause urothelial cells to release more ATP in the sub-urethral space, potentially leading to mitochondrial dysfunction and cell apoptosis.

Commensal bacteria are more abundant than pathogens in the bladder of healthy women and are associated with a reduced risk of UUI by inhibiting the pathogenesis process [9]. We surmised that they might have a protective role against extracellularly deposited bacterial ATP. This was supported by experiments showing that *L. crispatus* and *L. gasseri* did not release significant amounts of ATP [Fig 3B], and *L. crispatus* could reduce ATP levels in AU supplemented with ATP 0.1 mM [Fig 3C]. In addition, *L. crispatus* and *L. gasseri* inhibited the stimulation of calcium influx caused by *E. coli*-derived compounds [Fig 2A/B and C]. Preliminary evidence was obtained that commensal bacteria could degrade or utilize ATP, with *L. crispatus* reducing its levels in AU. *Lactobacillus crispatus* also increased *MAOB* gene expression [Fig 5B], encoding enzymes that can degrade biogenic amines neuroactive chemicals. A decrease in the level of these mitochondrial enzymes has been postulated to worsen neurological disorders and may also be another mechanism by which commensal bacteria mitigate the effects of these chemicals [17].

Lactobacilli are typically restricted to glycolytic and fermentative pathways which produce much less ATP than through the respiratory pathways used by other bacteria. If lactobacilli present in the bladder microbiota or even the vagina, can scavenge ATP it may not only potentially provide an extra energy source for the bacteria, but could sequester it away from the epithelial layer thereby promoting a homeostatic environment. These are important findings, since ATP promoted collagen matrix contraction by myofibroblasts [Fig 7A/D], an *in vitro* model of smooth muscle contraction, suggesting a mechanism for premature voiding and the potential for lactobacilli strains to interfere with this process. However, not all strains of lactobacilli tested were protective against the effects of ATP. *Lactobacillus vaginalis,* commonly found in the oral, vaginal, and intestinal microbiomes, has been associated with intermediate grades of bacterial vaginosis [18]. In this study, *L. vaginalis* was found to release 0.314 ± 0.023 uM ATP [Fig 3B], several fold more than *E. coli,* which suggests that certain lactobacilli may in fact be part of the disease process, though will require more investigation.

The neurotransmitter GABA is produced by bacteria [19] including certain species of *Lactobacillus*, and we showed that while it did not cause contraction of myofibroblasts [Fig 7], it could inhibit contraction caused by *E. coli* [Fig 7E]. The increase in intracellular calcium levels results in the secretion of ATP by urothelial cells [Fig 3K], with two potential mechanisms likely. ATP can be released via channels, such as the connexin hemichannels, pannexin as well as several anion channels [20]. It is possible that stimulation of calcium influx in urothelial cells may cause increased expression of vesicular nucleotide transporter (VNUT) in the cell and subsequent release of ATP into the sub-urethral and muscle layer causing bladder contraction. The alternative is for a continuously activated calcium channel leading to mitochondrial calcium overload, apoptosis and release of ATP from urothelial cells [21].

Alpha smooth muscle actin (α-SMA) has a well substantiated, central role in the production of contractile force during wound healing and fibro-constrictive diseases [22]. Confocal and qPCR results herein show a direct correlation between increased α-SMA immunoreactivity and uropathogen induced contraction of the collagen gel matrix by myofibroblasts *in vitro* [Fig 8B/C and 7B]. There was also a correlation between decreased α-SMA immunoreactivity and a decrease in collagen matrix contraction induced by *L. crispatus* [Fig 8B/C and 7B]. Increased intracellular calcium levels can drive the urothelial cells to the apoptosis phase. TNF-alpha can be an inducer of apoptosis [23], and so the ability of *L. crispatus* to reduce the *E. coli*-stimulated upregulation of this gene in myofibroblast cells, could be significant [Fig 8D].

In summary, we report the discovery of a potential means by which commensal members of the urinary microbiota, in particular *L. crispatus* and *L. gasseri*, can mitigate the ability of uropathogenic *E. coli* to stimulate pathways associated with conditions such as UUI.

## MATERIALS and METHODS

### Bacterial supernatant preparation

*Escherichia coli* 1A2 UPEC was maintained on LB agar (Difco, MD), *Lactobacillus gasseri* KE-1 (urinary isolate)*, Lactobacillus crispatus* ATCC 33820, *Enterococcus faecalis* ATCC 33186, were maintained on MRS agar (Difco, MD), *Gardnerella vaginalis* ATCC 14018, *Lactobacillus vaginalis* NCFB 2810 were maintained on CBA and *Gardnerella* Selective Agar. For these studies, all strains of bacteria were grown in artificial urine [24] which in preliminary experiments was shown not to stimulate the influx of calcium when in the presence of human cell lines.

Supernatants were collected from cultures grown overnight (24 hours) at 37°C after reaching stationary phase. Cultures were pelleted by centrifugation at 5000 rpm (Eppendorf Centrifuge 5804 R) for 15 minutes. The supernatant was pH adjusted to 7.0 with 0.1 Molar HCL or NaOH, filter sterilized with 0.22 um sterile syringe filter, and aliquoted and stored at −20 C° until use. In the case of *E. coli* and *E. faecalis*, overnight cultures were diluted 1:100 with fresh artificial urine, returned to incubation at 37 °C and sampled at T= 1, 2, 3, 4, 5 and 24 hours for testing. For the experiments involving the addition of supernatants from *L. crispatus* or *L. gasseri* to that from uropathogens, the urothelial cells were first treated with *L. crispatus* or *L. gasseri* supernatant for one minute, then the uropathogenic supernatant was added. In the case of serial dilution, *L. crispatus* supernatant was diluted for 6-fold to the *E. coli* supernatant.

For investigation the subtherapeutic concentration of ciprofloxacin, the *L. crispatus* was grown in deMan, Rogosa, Sharpe media (MRS, Difco, MD). Growth curves for these bacteria were generated using a plate reader (Eon Biotek, VT) at OD600 and 37°C to determine exponential phase.

### Cell culture

Bladder epithelial cells (5637 - ATCC HTB-9) were maintained in RPMI 1640 (Roswell-Park Memorial Institute media –; Thermo Fisher Scientific, MA) supplemented with 10% fetal bovine serum (FBS) (Thermo Fisher Scientific, Ma.) and 2 mM L-glutamine (Thermo Fisher Scientific, MA.) at 37°C and 5% CO_2_. The media was changed every 48 hours or more regularly if the cells were confluent (90%-100%), after washing by 1X PBS and trypsinization by 0.25% Trypsin-EDTA (1X) (Gibco), with the ratio of 1 to 10. Primary myofibroblast cells were extracted from the palmar fascia during surgery from normal tissue. Primary cultures were maintained in DMEM with 10% fetal bovine serum (FBS; Life Technologies, Carlsbad, CA, USA), 1% L-glutamine (Life Technologies) and 1% antibiotic-antimycotic solution (Life Technologies) at 37°C in 5% CO2. All primary cell lines were used up to a maximum of four passages, after which they were discarded.

### RNA isolation and qPCR from cell lines

RNA was isolated from the samples (200 ng/uL) using the Ambion by Life Technologies Purelink™ RNA mini kit (Thermo Fisher Scientific, MA), following the manufacturer’s instructions. cDNA was made following the instructions on the Applied Biosystems High Capacity cDNA Reverse Transcription Kit (Thermo-Fisher Scientific, MA) and PCR was conducted using a Master Cycler gradient PCR thermal cycler (Eppendorf, NY). Using GAPDH as the housekeeping gene, qPCR was set up with each sample being run on the plate in triplicate for each of the conditions. A list of the primer sequences used can be found in Table 1. Power SYBR Green PCR Master Mix was used (Thermo Fisher Scientific, MA).

### Fluorescent microscopy of calcium influx of 5637 cells

The influx of calcium was measured using the Fluo-4 DirectTM Calcium Assay kit (InvitrogenTM, CA). Samples and reagents were prepared according to the protocol manual provided. Ninety-six well plates were seeded with 100 μl of 5637 cells at 1×10^5^ cells/mL in supplemented RPMI and allowed to reach confluency, which occurred at about 48-72 hours. Cells were counted by using the Invitrogen Countess Automated Cell Counter (Thermo Fisher Scientific, MA.) per the manufacturers’ instructions. Fifty microliters of cell culture media were removed from the initial 100 μl and 50 μl of Fluo-4 DirectTM calcium reagent was added to each well. The plate was incubated at 37° C for 30 minutes at room temperature while protected from light. Controls included ionomycin (1 uM, Sigma ≥98% HPLC), ATP (1 uM, Sigma A1852), GABA (1uM, Sigma BioXtra ≥99%) and LPS (0.13 milligram/mL, Sigma L3755). The effect of treatments was assessed using a Nikon epifluorescence Ts2R scope at 10x magnification at 494nm for excitation and 516 nm for emission for 60 seconds. The image intensity was calculated using ImageJ and is indicative of Ca2+ influx into the urothelial cell’s cytoplasmic space from either the extracellular environment or intracellular Ca2+ stores (here on out just referred to as Ca2+ influx).

### Quantification of ATP

A luminescent assay kit (BacTiter-Glo™ Microbial Cell Viability Assay, G8230) was used to quantify the amount of extracellular ATP released by the bacteria into the supernatant and released by the cells into the cell media. The Synergy™ H4 Hybrid Multi-Mode Microplate Reader was used to quantify the amount of extracellular ATP.

### Myofibroblast populated collagen contraction

A collagen matrix was set up using 1.8 mg/ml sterile collagen and a neutralization solution [25]. The neutralization solution was made by mixing Waymouth Media (Sigma, W1625) and 2 parts 0.34M NaOH (Sigma, 221465). One-part neutralization mixture was then added to 4 parts collagen, mixed with 1×10^5^ cells to a final volume of 500 μl and added to each well in a 24 well plate. After 45-minute incubation at 37°C, 1 mL 2% FBS was added to each well and the plate was incubated for an additional 72 hours at 37 °C. The media was then removed, fresh media and treatment was added, and the collagen matrix was released using a sterile spatula. The plate was scanned using a Canon PIXMA MP250 immediately after release and also at 1, 3, 5 and 24 hours. The size of the collagen matrix was measured using ImageJ and the percent contraction was calculated. To decrease any shock to the myofibroblast, all bacterial strains were grown in DMEM with 2% FBS.

### Immunocytochemistry

Myofibroblast cells were cultured in a μ-Slide 8 Well (ibidi, 80826) to become fully confluent (90%-100%). Cell were fixed with paraformaldehyde for 10 minutes at room temperature, then permeabilized with 0.1% Triton X-100 in PBS. Non-specific staining was blocked with Background Sniper (Biocare Medical, BS966). Cells were stained by incubating with the monoclonal anti-actin, α-smooth muscle (Sigma, A2547) diluted 1∶200 and using Alexa Fluor 488 Donkey anti-mouse IgG secondary antibody (ThermoFisher, A-21202) to detect the fluorescence. The cells were washed, excess liquid aspirated, and secondary antibody solution was added (1-10ug/ml) (Alexa Fluor 488 Donkey anti-mouse IgG secondary antibody, ThermoFisher, A-21202). DAPI staining was used for nuclei. Confocal images were obtained with a Nikon Eclipse Ti2 (X60 objective lens, Nikon, Canada). Fluorescence intensity measurements were obtained from entire cells and analyzed with Image J software. Control specimens were identical to experimental specimens except they were exposed to irrelevant isotype matched antibody.

### Myofibroblast populated collagen RNA extraction and qPCR

After incubation and aspiration of media, the collagen matrix was collected in microcentrifuge tubes for high speed centrifugation for 5 minutes and then the supernatant was discarded. An aliquot of 100 uL pre-warmed 0.25 mg/ml collagenase was added to each tube and incubated for 15 minutes at 37°C. RNA was isolated from the samples using the Direct-zol RNA Miniprep Kit (Zymo Research) following the manufacturer’s instructions, and Trizol reagent was used to lyse the samples. The RNA concentration was measured using nanodrop. cDNA was made following the instructions on the Applied Biosystems High Capacity cDNA Reverse Transcription Kit (Thermo-Fisher Scientific, MA) and PCR was conducted using a MasterCycler gradient PCR thermal cycler (Eppendorf, NY). Quantitative PCR was set up with each sample being run on the plate in triplicate for each of the conditions, as described earlier. GAPDH was also used as the housekeeping gene, A list of the primers used can be found in Supplementary Table 1.

### Statistics

The data are expressed as mean ± SEM. Statistical significance was assessed using one-way ANOVA followed by Tukey’s test (GraphPad Prism 5)

## Acknowledgments

This project was funded by Kimberly Clark Corporation who were involved in the study design, analysis and preparation of the manuscript.

---

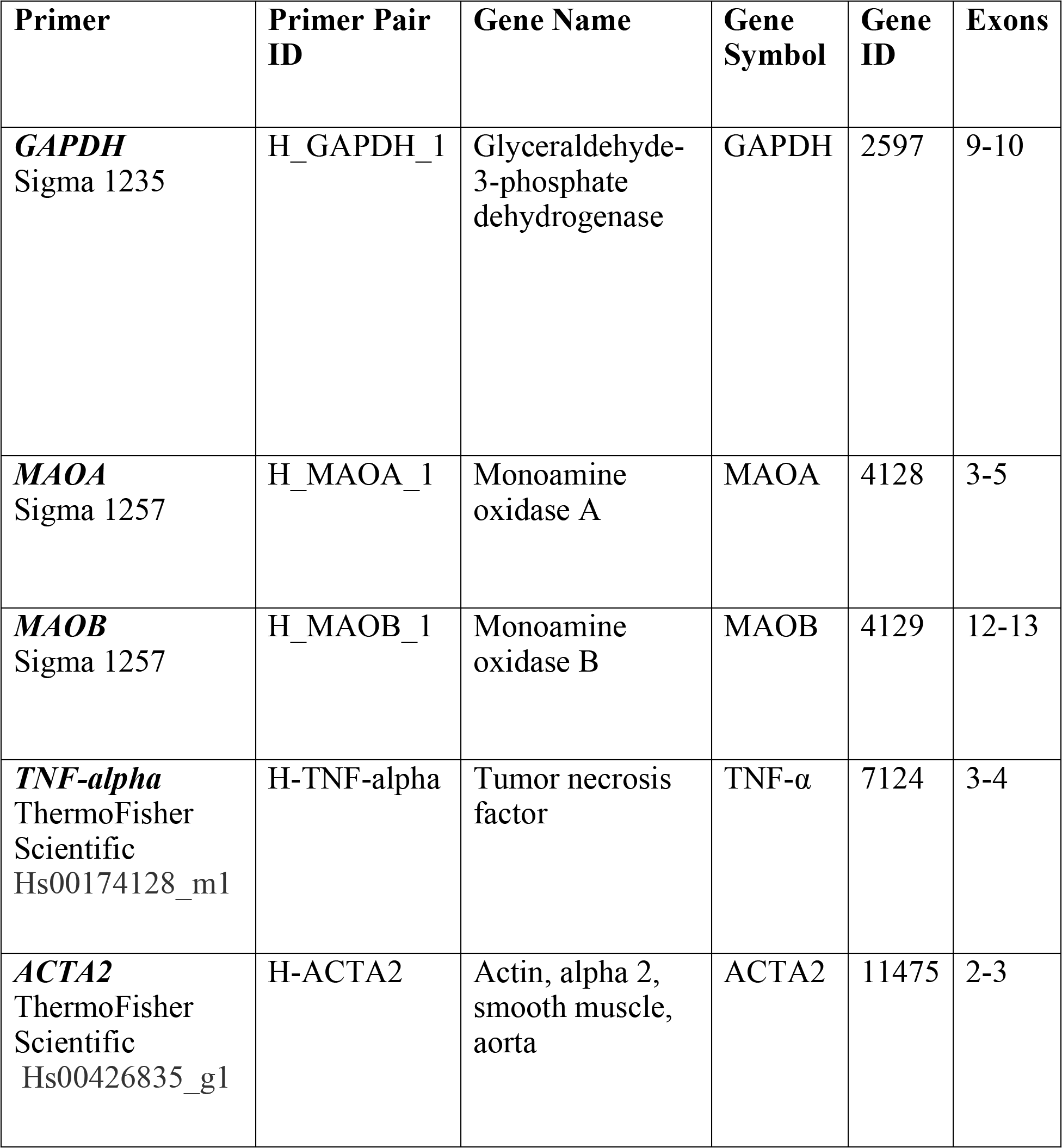
Supplementary Table 1.

